# Endoplasmic Reticulum Retention Motif Fused to Recombinant Anti-cancer Monoclonal Antibody (mAb) CO17-1A Affects mAb Expression and Plant Stress Response

**DOI:** 10.1101/335646

**Authors:** Ilchan Song, Yang Joo Kang, Young Koung Lee, Soon-chul Myung, Kisung Ko

**Affiliations:** Department of Pathology, College of Medicine, Chung-Ang University, Seoul, Korea; Department of Medicine, College of Medicine, Chung-Ang University, Seoul, Korea; Cold Spring Harbor Laboratory, Cold Spring Harbor, NY 11724, USA; Department of Urology, College of Medicine, Chung-Ang University, Seoul, Korea; Division of Biological Science and Research Institute for Glycoscience, Wonkwang University, Iksan, Korea

## Abstract

The endoplasmic reticulum (ER) is the main site of protein synthesis, folding, and secretion to other organelles. The capacity of the ER to process proteins is limited, and excessive accumulation of unfolded and misfolded proteins can induce ER stress, which is associated with plant diseases. Here, a transgenic *Arabidopsis* system was established to express anti-cancer monoclonal antibodies (mAbs) that recognize the tumor-associated antigen GA733-2. The ER retention Lys-Asp-Glu-Leu (KDEL) motif sequence was added to the C-terminus of the heavy chain to retain anti-colorectal cancer mAbs in the ER, consequently boosting mAb production. *Agrobacterium*-mediated floral dip transformation was used to generate T_1_ transformants, and homozygous T_4_ seeds obtained from transgenic *Arabidopsis* plants expressing anti-colorectal cancer mAbs were used to confirm the physiological effects of KDEL tagging. Germination rates were not significantly different between mAb CO and mAb COK. However, mAb COK primary root lengths were shorter than those of mAb CO plants and non-transgenic *Arabidopsis* plants in *in vitro* media. Most ER stress-related genes, with the exception of *bZIP28* and *IRE1a*, were upregulated in mAb COK plants compared to mAb CO plants. Western blot and SDS-PAGE analyses showed that mAb COK plants exhibited up to five-times higher expression and mAb amounts than mAb CO plants. Enhanced expression in mAb COK plants was confirmed by immunohistochemical analyses. mAb COK was distributed across most of the area of leaf tissues, whereas mAb CO was mainly distributed in extracellular areas. Surface plasmon resonance analyses revealed that both mAb CO and mAb COK possessed equivalent or slightly better binding activities to antigen EpCAM compared to a commercially available parental antibody. These results suggest that the introduction of the KDEL motif is a promising strategy for obtaining enhanced amounts of recombinant therapeutic proteins, but the KDEL sequence may induce ER stress and slightly reduce plant biomass.

## Introduction

Colorectal cancer (CRC) is the second most frequently diagnosed cancer in women and the third most commonly diagnosed cancer in men worldwide [1]. Highly valuable recombinant therapeutic antibodies for the diagnosis and treatment of cancer have been produced in transgenic plant systems [2, 3]. In fact, to express anti-CRC mAbs, *Arabidopsis* was selected as a production platform because of its short life cycle and high total soluble protein content [4, 5]; however, therapeutic glycoproteins produced in plant cells had plant-specific *N*-glycans. The fusion of the endoplasmic reticulum (ER) retention KDEL (Lys-Asp-Glu-Leu) motif is frequently used to retain and accumulate recombinant therapeutic proteins in the ER, enhancing their stability and production yields in plants. KDEL is added to the C-terminus of the heavy chain (HC) to induce high mannose glycan structures without plant-specific glycan residues [α(1,3)-fucose and β(1,2)-xylose], which can trigger a host immune response [6, 7].

The accumulation of recombinant anti-colorectal cancer mAbs in the ER requires a large ER-mediated protein quality control (ERQC) capacity, which results in ER stress [8-10]. ER stress activates three main unfolded protein response (UPR) signaling pathways [11, 12], namely ER-membrane-associated activating transcription factor 6 (ATF6), inositol-requiring enzyme 1 (IRE1), and protein kinase RNA-like ER kinase (PERK) [13, 14]. In this study, three basic-leucine zipper transcription factor family proteins (bZIP17, bZIP28, and bZIP60), two binding immunoglobulin proteins (BiP1 and BiP3), two plant-specific NAC transcription factors (NAC103 and NAC089), regulator of ER stress-induced programmed cell death (BAX inhibitor 1), B-cell lymphoma 2 (Bcl-2)-associated athanogene 7 (BAG7), and ER oxidoreductin 1 (ERo1) were analyzed in terms of ER stress regulation [15-17].

We obtained homozygous seeds from transgenic *Arabidopsis thaliana* plants expressing anti-CRC mAb^P^s (mAb CO and mAb CO tagged with KDEL (mAb COK)) to recognize CRC-associated antigen GA733 (EpCAM), which is highly expressed in CRC cells [18, 19]. To confirm the effects of ER retention motif tagging on transgenic *Arabidopsis*, the KDEL motif was fused to the C-terminus of the original HC. Germination rates, primary root lengths, regulation of ER stress-related genes, transcriptional and translational levels, subcellular distribution of mAbs in fresh leaves, purification amount per unit fresh leaf, and antigen-binding functions were compared between the non KDEL-tagged group (mAb CO) and the KDEL-tagged group (mAb COK).

## Materials and Methods

### Vector construction and *Arabidopsis* transformation

Plant expression vectors pBI CO17-1A (pBI CO) and pBI CO17-1AK (pBI COK), carrying mAb CO and mAb CO tagged with the KDEL ER retention motif (mAb COK), were transformed into *Agrobacterium tumefaciens* strain GV3101::pMP90 via electroporation. Wild-type *Arabidopsis* plants were transformed using the floral dip method [20] (Fig 1). Approximately one month prior to transformation, non-transgenic (NT) *Arabidopsis* seeds were sown with approximately 4–5 plants per pot in eight pots. The plants were grown under standard conditions (16 h light/8 h dark) in a growth chamber. To induce the proliferation of multiple secondary bolts, the first bolts of *Arabidopsis* were trimmed. Two-days before plant transformation, *A. tumefaciens* strain GV3101::pMP90, carrying mAb CO or mAb COK expression cassettes, was cultured at 28–30°C in LB (Luria-Bertani) with kanamycin. *Agrobacteria* were centrifuged (4,000 rpm for 10 min) and pellets were resuspended with infiltration media until the OD values were adjusted to 0.8–1.0. Before transformation, Silwet L-77 was added to the infiltration media at a concentration of 0.02%. The pots were inverted into the infiltration solution for 5 min. After dipping transformation, pots were covered with a black plastic bag to ensure highly humid and dark conditions, and the trays were placed in a growth chamber. The following day, the plastic bags were removed, and the plants were maintained under standard conditions in a growth chamber until seeds were well-ripened. Transformed seedlings were selected on agar plates containing Murashige and Skoog medium (adjusted pH 5.7) (10 g·L^−1^ of sucrose, 8 g·L^−1^ of plant agar, and 4.3 g·L^−1^ of MS B5 vitamin; Duchefa Biochemie, Haarlem, Netherlands), containing 50 mg·L^−1^ kanamycin and 25 mg·L^−1^ cefotaxime. All transformed seedlings were grown in a growth chamber at 22°C under a 16 h light/8 h dark cycle. After T_1_ seeds were obtained, kanamycin selection was repeated over generations until homozygous lines were obtained. Homozygous T_4_ seeds were used in this experiment.

**Fig 1.**
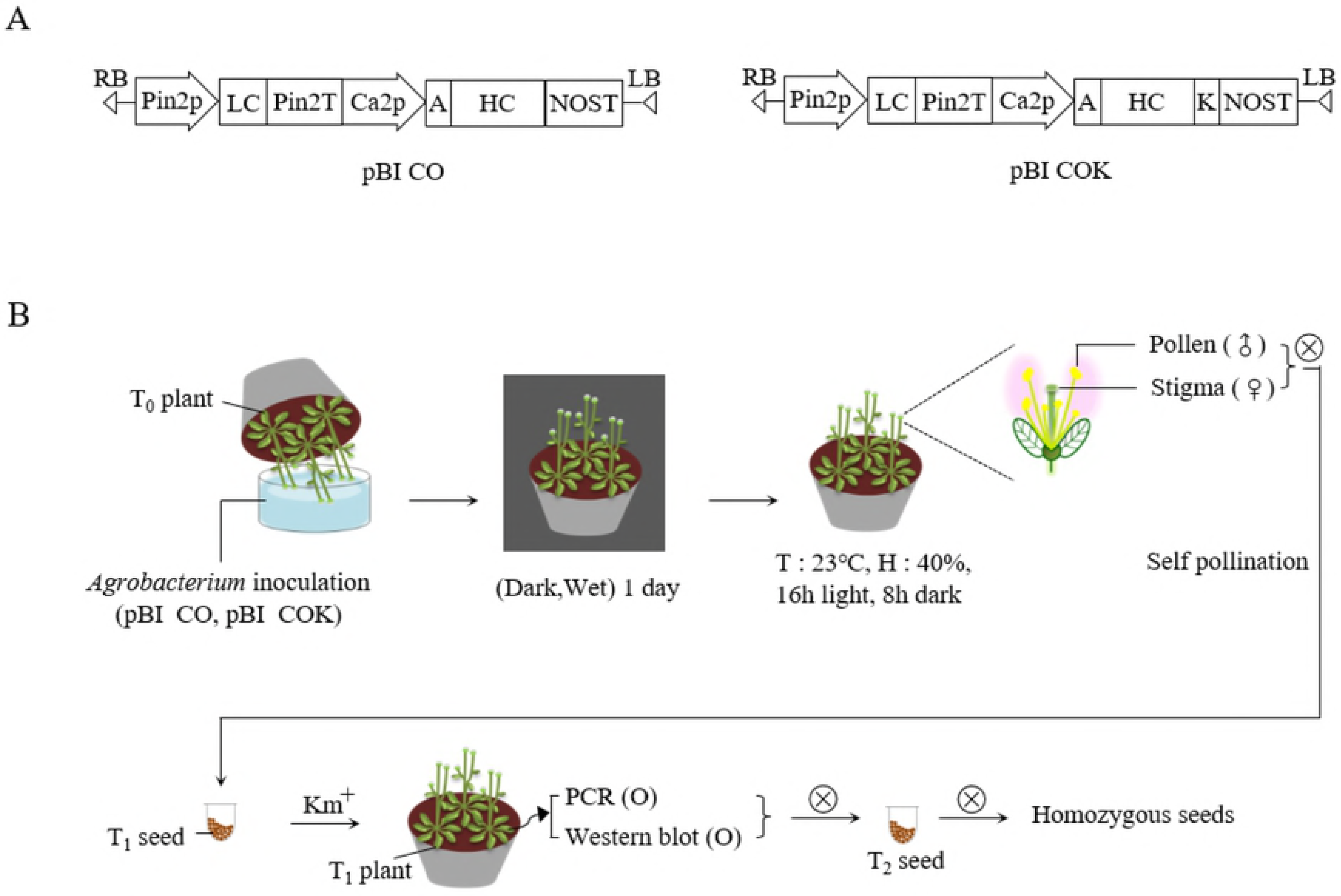
Schematic diagram of the floral dip transformation process. *Agrobacterium tumefaciens* strain GV3101::pMP90, carrying plant binary vectors pBI CO and pBI COK, were used for floral dip transformation. (A) Schematic diagram of mAb CO and mAb COK gene expression cassettes in plant expression vector pBI121, which was used for *Agrobacterium*-mediated transformation. Pin2p, promoter of the *Pin2* gene from potatoes, and Ca2p, the cauliflower mosaic virus 35S promoter, control the light and heavy chains, respectively. KDEL is the 3′ endoplasmic reticulum (ER) retention motif. (A) An alfalfa mosaic virus untranslated leader sequence of RNA4; Pin2T, terminator of the *Pin2* gene from potato; NOST, terminator of nopaline synthase (*NOS*) gene. (B) After floral dip transformation, plants were placed in a box, watered, and maintained under a dark and humid atmosphere for one day. Plants were cultured for a few weeks until mature seeds were obtained. Seeds were germinated on kanamycin-containing medium to select transformants. Kanamycin selection was repeated over generations until homozygous lines were identified.

### PCR analysis

Rosette leaves (approximately 100 mg) from four-week-old NT and transgenic plants expressing mAb CO were used for polymerase chain reaction (PCR) analysis. A DNA extraction kit (RBC Bioscience, Seoul, Korea) was used to extract genomic DNA from plant leaves, following the manufacturer’s recommended protocol. PCR was performed to confirm the presence HC (1,416 bp) and LC (717 bp) genes associated with mAb CO, and transformants were selected from T_1_ plants. Primers were designed as follows: HC forward primer, 5′-GCGAATTCATGGAATGGAGCAGAGTCTT TATC-3′; HC reverse primer, 5′-GATTAATCGATTTTACCCGGAGTCCG-3′; LC forward primer, 5′-GCCTCGAGATGGGCATCAAGATGGAATCACAG-3′; LC reverse primer, 5′ GAGGTACCCTAACACTCATTCCTGTTGAAGCTC-3′. Leaves from NT plants were used as a negative control, and the pBI CO gene was used as a positive control. PCR analysis was replicated more than three times.

### Western blot analysis

The rosette leaves of T_1_ plants, with confirmed foreign gene insertion via PCR, were subjected to western blot analyses to confirm target protein expression. Fifty milligram of fresh rosette leaves was frozen in liquid nitrogen and immediately crushed. To extract total soluble protein, leaf samples were mixed with 100 μL of sample buffer (1 M Tris-HCl, 50% glycerol, 10% SDS, 5% 2-mercaptoethanol, and 0.1% bromophenol blue), boiled for 10 min, and cooled on ice. Total soluble proteins were separated by 12.5% SDS-PAGE and transferred to a nitrocellulose membrane (Millipore, Billerica, MA). Membranes were incubated with 5% skimmed milk (Sigma, St. Louis, MO) for 16 h at 4°C. The nitrocellulose blots were incubated with goat anti-murine IgG Fcγ and anti-murine IgG F(ab)′2, which recognize HC and LC of mAb CO, respectively. After washing three times for 10 min at room temperature, proteins were detected with the SuperSignal West Pico Chemiluminescent Substrate (Thermo Scientific, Rockford, IL), and X-ray film (Fuji, Tokyo, Japan) was used for visualization. Rosette leaves of NT *Arabidopsis* plants were used as a negative control.

### Plant growth conditions and morphological assessments

After three continuous screenings on kanamycin media, homozygous T_4_ seeds were obtained. Homozygous T_4_ seeds of transgenic *Arabidopsis* plants expressing mAb CO were used as the material for this experiment. Two-hundred surface-sterilized seeds of each plant expressing mAb CO (CO) and mAb COK (COK), including NT plants, were sown on agar plates containing Murashige and Skoog medium (adjusted pH 5.7; 10 g·L^−1^ of sucrose, 8 g·L^−1^ of plant agar, and 4.3 g·L^−1^ of MS B5 vitamin (Duchefa Biochemie, Haarlem, Netherlands)), containing 50 mg·L^−1^ kanamycin and 25 mg·L^−1^ cefotaxime. The seedling plates were vernalized at 4°C for two days under dark conditions to improve the rate and synchrony of germination, and were then placed vertically in a growth chamber at 23°C under 16 h light/8 h dark conditions. The germination rates were calculated as follows: (number of germinated seeds/total seeds) ×; 100. Germination was checked four days after transfer to the growth chamber, and primary root lengths were checked at four, six, eight, 10, and 12 days. Thirty-two shoots of each plant (CO, COK, and NT) were transferred to pots for further study. Rosette leaf lengths were measured from the petiole to the leaf blade using a ruler (in cm) over a period of four weeks at one-week intervals. After 10 weeks of soil culture, the roots of 96 plants were rinsed with tap water for 1 min, and the main root lengths were measured using a ruler. Photographs were taken with a camera (Digital Gross System) (Humintec, Suwon, Korea), and at least two independent experiments were conducted for all data analyses.

### RNA extraction and reverse transcription (RT) PCR analyses

Total RNA was extracted from NT plants and transgenic *Arabidopsis* plants expressing mAb CO and mAb COK that were grown under standard conditions in a growth chamber. Fresh rosettes leaves were sampled and homogenized after freezing in liquid nitrogen. RNA was extracted using TRIzol, as previously described [21, 22]. Extracted RNA was stored in a deep freezer (−70°C) for further study. Genomic DNA was removed, and cDNA was synthesized using a Quantitect reverse transcription kit (Quagen, Valencia, CA) according to the manufacturer’s instructions. Each cDNA sample was used as a template for RT-PCR analyses. RT-PCR was performed using the Maxime PCR Premix Kit (Intron Biotechnology, Seoul, Korea). Transcription of HC and LC genes was confirmed with primers as described above. Actin 8 was used as a housekeeping control gene, and the primer sets were used to amplify the gene: actin 8 forward primer, 5′-CAACTATGTTCTCAGGTATTGCAGA-3′; actin 8 reverse primer: 5′-GTCATGGAAACGATGTCTCTTTAGT-3′. NT plants were used as a negative control. RT-PCR analyses were performed in duplicate, and the reverse transcription PCR product was run on an agarose gel for UV detection.

### RNA isolation and quantitative real time PCR (qRT-PCR) analysis

*Arabidopsis* plants were cultivated (vertical growth) in MS (Murashige and Skoog) media for 14 days after seeding, and were then harvested to dissect root tissues. Total RNA was extracted from *Arabidopsis* roots using an RNeasy Mini Kit (Qiagen, Valencia, CA), and the extracted RNA samples were the processed with an RNase-free DNase kit (Qiagen, Valencia, CA), according to the supplied protocol. cDNA was synthesized from 2 μg of total RNA using the Superscript III First Strand Synthesis kit (Invitrogen, Carlsbad, CA). Regarding real-time qPCR, each primer was designed to amplify a 200–300 bp region of the corresponding transcript, and the control transcript for qRT-PCR was *UBI10*. Real-time PCR analyses were performed on a C1000 Thermal Cycler (Bio-Rad, Hercules, CA). Primers were tested on NT plants via quantitative PCR using the Bio-Rad CFX96 real time system, and primer quality was determined based on melting curve analyses and amplification efficiency. Gene expression was measured in three technical replicates for each sample. cDNA was generated via reverse transcription of 2 μg total RNA as described above. Primers were designed by AtRTPrimer [23], and the primer sequences used for qRT-PCR are provided in Table 1.

**Table 1.**
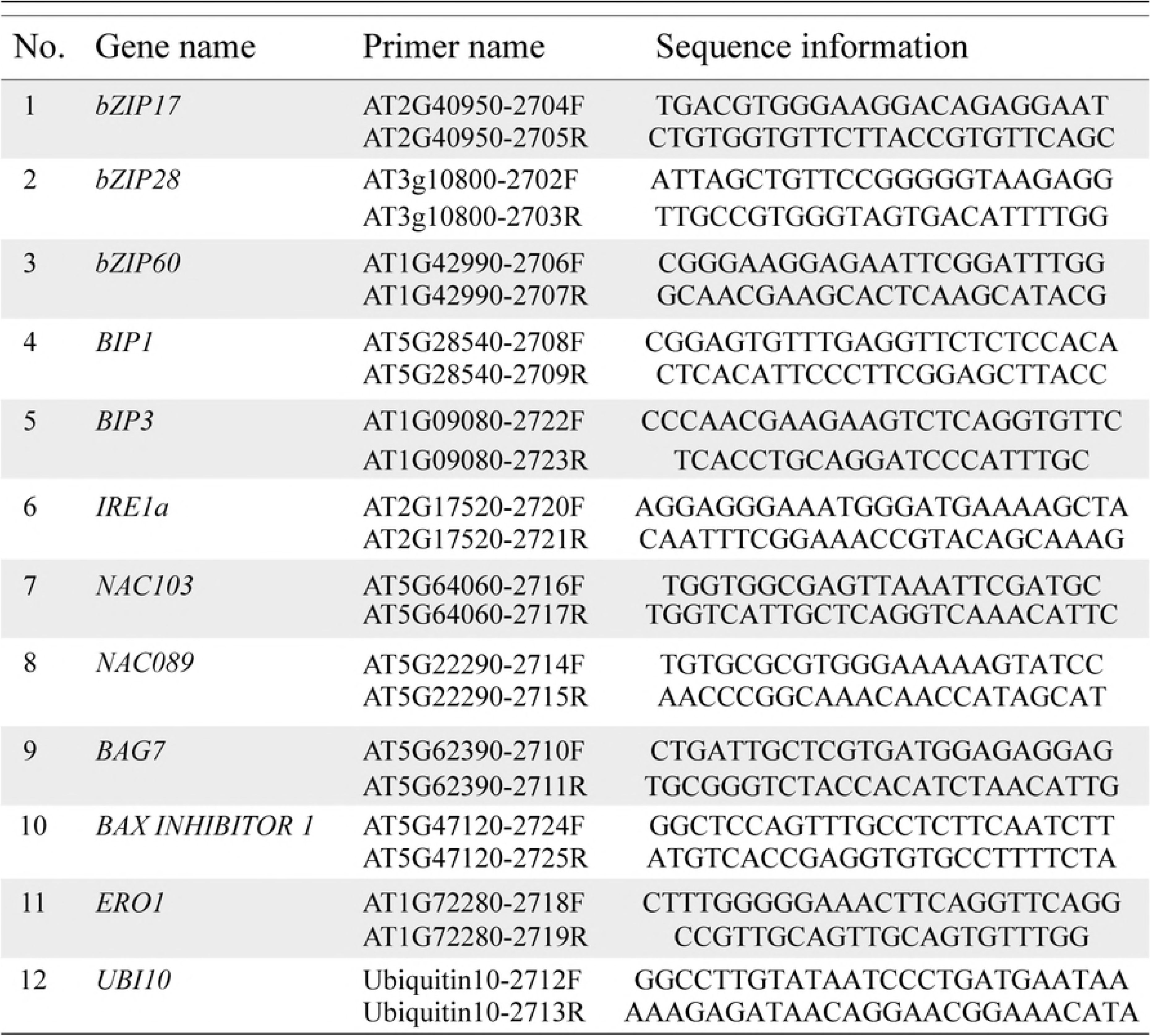
Primers specific for ER stress-related genes (*bZIP17, bZIP28, bZIP60, BiP1, BiP3, IRE1a, NAC103, NAC089, BAG7, BAX inhibitor 1*, and *ERO1*) in *Arabidopsis thaliana* (Han and Kim, 2006).

### Immunohistochemical analysis

The intracellular distributions of mAb CO and mAb COK in the rosette leaf tissues of transgenic *Arabidopsis* were assessed using immunohistochemical analyses. Rosette leaf tissues, freshly harvested from 3–4-week-old *Arabidopsis* plants, were fixed in 10% formalin overnight and processed for paraffin embedding. Paraffin sections were cut to 4 μm thickness by a microtome. The sections were deparaffinized and rehydrated through xylene and serially diluted ethanol. Slides were then treated with 3% H_2_O_2_ for 10 min to block endogenous peroxidase activity. After washing twice with 1×; PBS, tissue sections were treated with antigen retrieval solution (Dako, Glostrup, Denmark) and kept at 120°C for 5 min in a pressure cooker. Tissue sections were then cooled to room temperature (23°C). After treatment with a protein block serum-free reagent for 30 min, the Dako REAL^™^ EnVision^™^ Detection System (Dako, Glostrup, Denmark) was applied according to manufacturer’s instructions, and samples were then counterstained with Mayer’s hematoxylin (Muto Pure Chemicals CO., Tokyo, Japan). Slides were mounted with cover glass using Permount solution and visualized with a microscope (×;400 magnification, BX53F (Olympus, Tokyo, Japan)).

### Purification of anti-cancer mAb

To purify anti-CRC mAbs (CO and COK) from plants, 200 g of each transgenic *Arabidopsis* leaf was homogenized in a 1.2 L extraction buffer (37.5 mM Tris-HCl pH 7.5, 50 mM NaCl, 15 mM EDTA, 75 mM sodium citrate, and 0.2% sodium thiosulfate) using a HR2094 grinder (Philips, Seoul, Korea). After centrifugation at 8,800 ×; *g* for 30 min at 4°C, the supernatant was filtered through Miracloth (Biosciences, La Jolla, CA), and the pH was adjusted to 5.1 with acetic acid. The solution was further centrifuged at 10,200 ×; *g* for 30 min. The supernatant was filtered through Miracloth and adjusted to 7.0 with the addition of 3M Tris-HCl, and ammonium sulfate was added up to an 8% concentration. After centrifugation at 8,800 ×; *g* for 30 min at 4°C, ammonium sulfate was added to the supernatant to reach a 24% concentration, and samples were incubated at 4°C overnight. The final solution was centrifuged at 4°C for 30 min, and the pellet was resuspended in one-twelfth of the starting volume of extraction buffer. The obtained solution was centrifuged at 10,200 ×; *g* for 30 min at 4°C. Plant-derived mAbs (mAb^P^ CO and mAb^P^ COK) were purified using protein A Sepharose 4 Fast Flow (GE Healthcare, Piscataway, NJ), according to the manufacturer’s recommendations. Both mAb^P^ CO and mAb^P^ COK proteins were dialyzed with 1×; PBS (pH 7.4). The protein concentration was determined via Nanodrop analysis (Biotek, Highland, VT), and the purified protein was visualized using SDS-PAGE. Purified mAb proteins were stored at −70°C for further study.

### Surface plasmon resonance analysis

To compare the binding affinities of anti-cancer mAbs^P^ (CO and COK) to the epithelial cell adhesion molecule (EpCAM). Surface plasmon resonance [24] analyses were performed using a ProteOn XPR36 instrument (Bio-Rad, Hercules, CA). The EpCAM antigen (R&D systems, Minneapolis, MN) was immobilized onto the surface of a GLC sensor chip and stabilized with PBS-T buffer (PBS buffer containing 0.05% v/v Tween-20). After stabilization, anti-EpCAM mAb (600 nM), mAb^P^ CO (600 nM), and mAb^P^ COK (600 nM) were applied to the sensor chip with a flow rate of 80 μl·min^−1^ at 25°C. The surface of the GLC chip was regenerated using phosphoric acid. Data analyses were performed with ProteOn Manager 2.1 software, and data were corrected by subtraction of the zero antibody concentration column as well as interspot correction.

## Results

### Generation of T_1_ *Arabidopsis* plants expressing anti-CRC mAb CO and mAb COK

Plant expression vectors pBI CO and pBI COK were transferred into *Arabidopsis* plants via *Agrobacterium-*mediated dipping transformation (Fig 1). A total of 2,000 T_1_ seeds were obtained after floral dip transformation, and seeds were then plated on MS medium containing kanamycin prior to transformant selection. Most of the sown seeds failed to develop true leaves, and were etiolated with light yellow-colored shoots (Fig 2A; top). Approximately 40 seedlings with true leaves that survived culture in the kanamycin medium, for each transgenic plant expressing mAb CO and mAb COK (CO and COK, respectively), were transplanted to a growth chambers under standard conditions (Fig 2A; bottom). The rosette leaves of 40 plants were used to confirm transgene insertion. PCR analysis of HC (1,416 bp) and LC (717 bp) gene products were observed in all *Arabidopsis* plants (Fig 2B; top). No PCR bands were observed in NT *Arabidopsis* (Fig 2B; top). After confirmation of transgene insertion, protein expression of HC and LC proteins in leaves was determined by western blot analysis (Fig 2B; bottom), and HC and LC protein bands were detected at approximately 50 and 25 kDa, respectively. Among the randomly selected 10 plants, six CO and five COK plants exhibited HC and LC protein bands, respectively (Fig 2B; bottom). Kanamycin selection was repeated across generations to confirm homozygous T_4_ seeds for plant growth observations and the stable mass production of transgenic *Arabidopsis* plants expressing mAb CO or mAb COK (data not shown). Three lines of CO plants and two lines of COK plants were obtained using repeated antibiotic selections (Fig S1).

**Fig 2.**
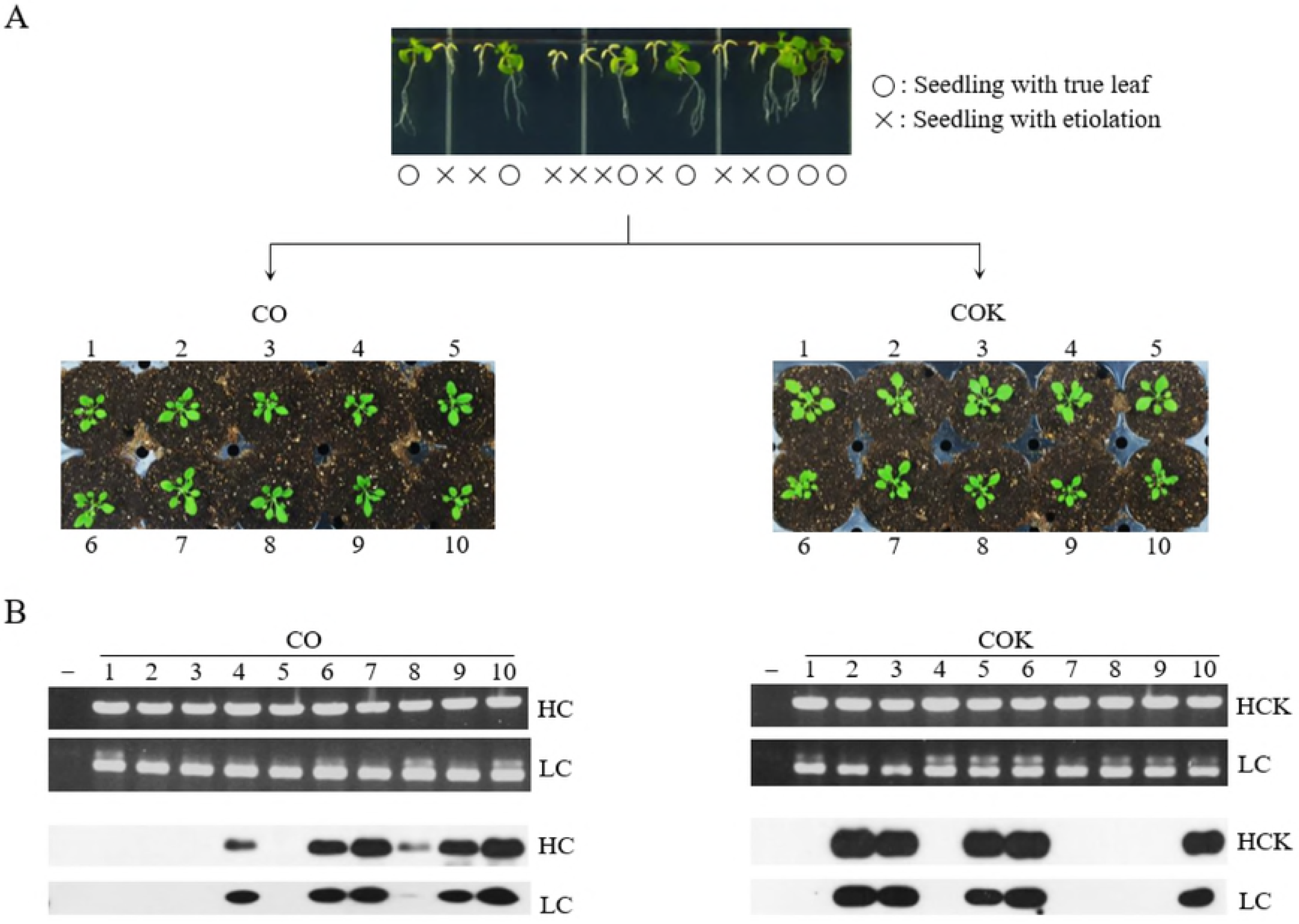
Screening T_1_ transformants expressing mAb CO and mAb COK using antibiotic selection, PCR, and western blot analyses. Selection of transformants after sowing T_1_ seeds on kanamycin media (top). o: seedling with true leaf; ×;: seedling with etiolation. Surviving shoots were transplanted to soil pots and placed in a growth chamber with 16 h of light and 8 h of darkness at 23°C (middle). Ten rosette leaves (mAb CO and mAb COK) were sampled from each T_1_ plant to confirm the existence of HC and LC gene insertions and protein expression levels (bottom). Goat anti-murine IgG Fcγ and goat anti-murine IgG F(ab)′2 were used to detect HC and LC, respectively.

### Effects of ER retention motif KDEL on germination rates and primary root growth of NT, CO, and COK seedlings in germination media with and without kanamycin

A total of 200 surface-sterilized seeds from each experimental group (NT, CO, and COK) were sown on MS agar plates containing kanamycin, and the same sets were sown on non-kanamycin media. In general, almost all seeds evenly started to germinate after two or three days of *in vitro* culture regardless of kanamycin presence. The germination rates of NT, CO, and COK in the non-kanamycin media were 98.6, 99.0, and 98.9%, respectively. The germination rates of NT, CO, and COK in the kanamycin-containing media were 94.7, 95.7, and 96.9%, respectively (Fig 3A). Twelve days after germination, the average primary root lengths of NT, CO, and COK specimens were 30.6, 28.0, and 20.7 mm, respectively, in media without kanamycin (Fig 3B; left). In kanamycin-containing media, the average primary root lengths of CO and COK were 23.2 and 12.2 mm, respectively, 12 days after germination (Fig 3B; right). In general, the primary root length of COK plants was shorter than that of plants in both the NT and CO groups (Fig 3B,C). All NT seeds were etiolated in MS media containing kanamycin within 4–5 days. The root lengths of plants in the CO group were approximately twice that of plants in the COK group (Fig 3C and S1 Fig). The primary root growth rate of COK was lower than that of CO in media, regardless of the presence of kanamycin.

**Fig 3.**
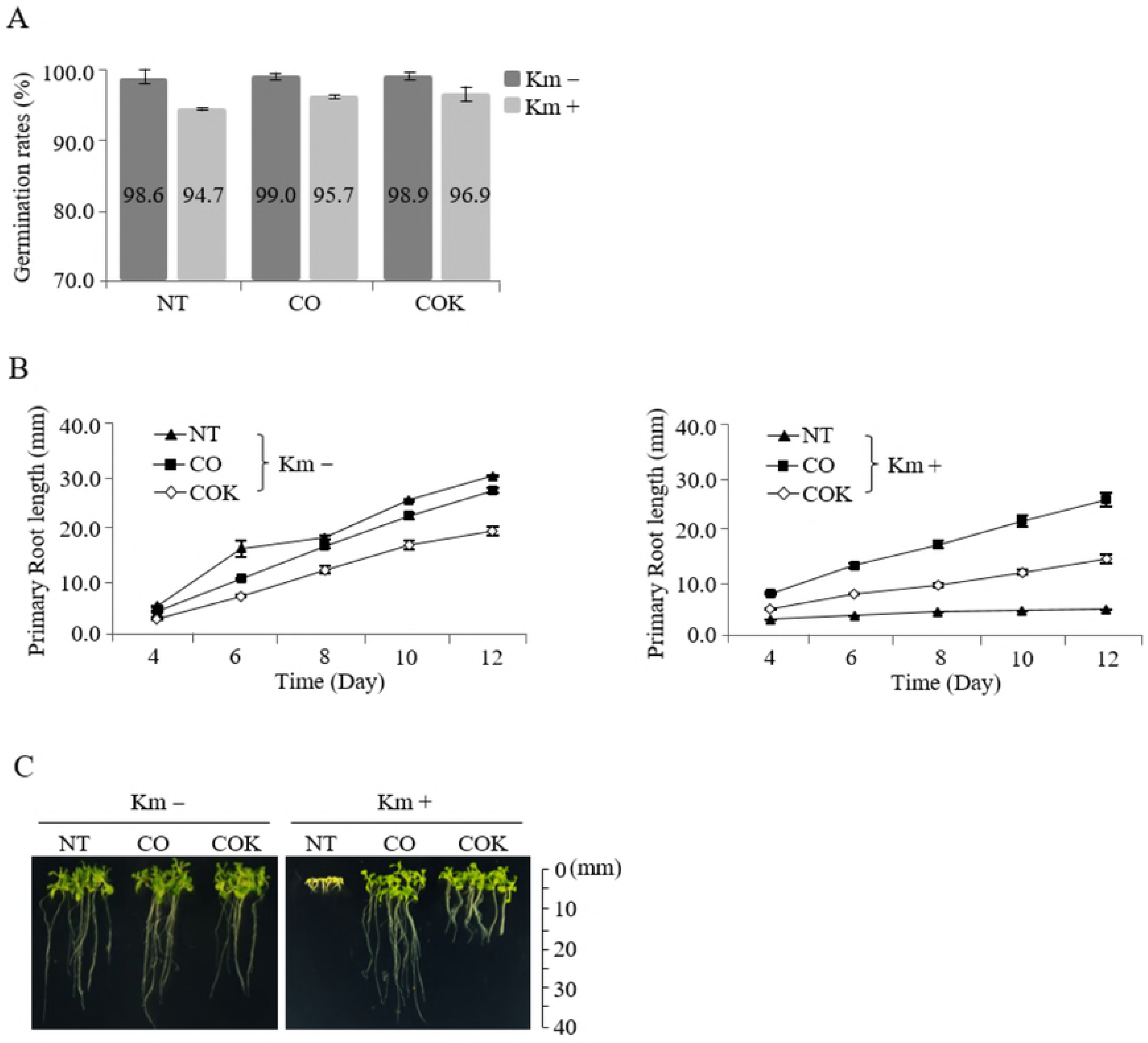
Effects of KDEL tagging on the plant growth of transgenic *Arabidopsis* plants expressing mAb CO and mAb COK under *in vitro* conditions with and without kanamycin. (A) Germination rates (number of germinated seeds/total seeds ×; 100) were calculated four days after the seeding of homozygous seeds on non-kanamycin or kanamycin-containing media. The values on the graphs represent the mean value of each experimental group. (B) Average primary root lengths of CO and COK plants *in vitro* with kanamycin (left) and without kanamycin (right) at four, six, eight, 10, and 12 days under the same conditions described above. (C) Photographs were taken 12 days after seeding on each MS media. Scales are displayed at 10 mm intervals. Standard deviation is indicated with error bars.

### mRNA expression of ER stress-related genes in transgenic *Arabidopsis* plants expressing mAb CO and mAb COK

The mRNA expression levels of 11 ER stress-related genes (*bZIP17, bZIP28, bZIP60, BiP1, BiP3, IRE1a, NAC103, NAC089, BAG7, BAX inhibitor 1*, and *ERO1*) were quantified using qRT-PCR analyses to investigate the effects of ER retention motif tagging on transgenic *Arabidopsis* plants expressing mAb CO and mAb COK (Fig 4). Expression of ER stress-related genes (*bZIP17, bZIP60, BiP1, BiP3, NAC103, NAC089, BAG7, BAX inhibitor 1*, and *ERO1*) were significantly upregulated in COK lines compared to CO lines, with the exception of *bZIP28* and *IRE1a* genes. Among the tested genes, the mRNA levels of *BiP3* in COK were almost three times higher compared to CO lines.

**Fig 4.**
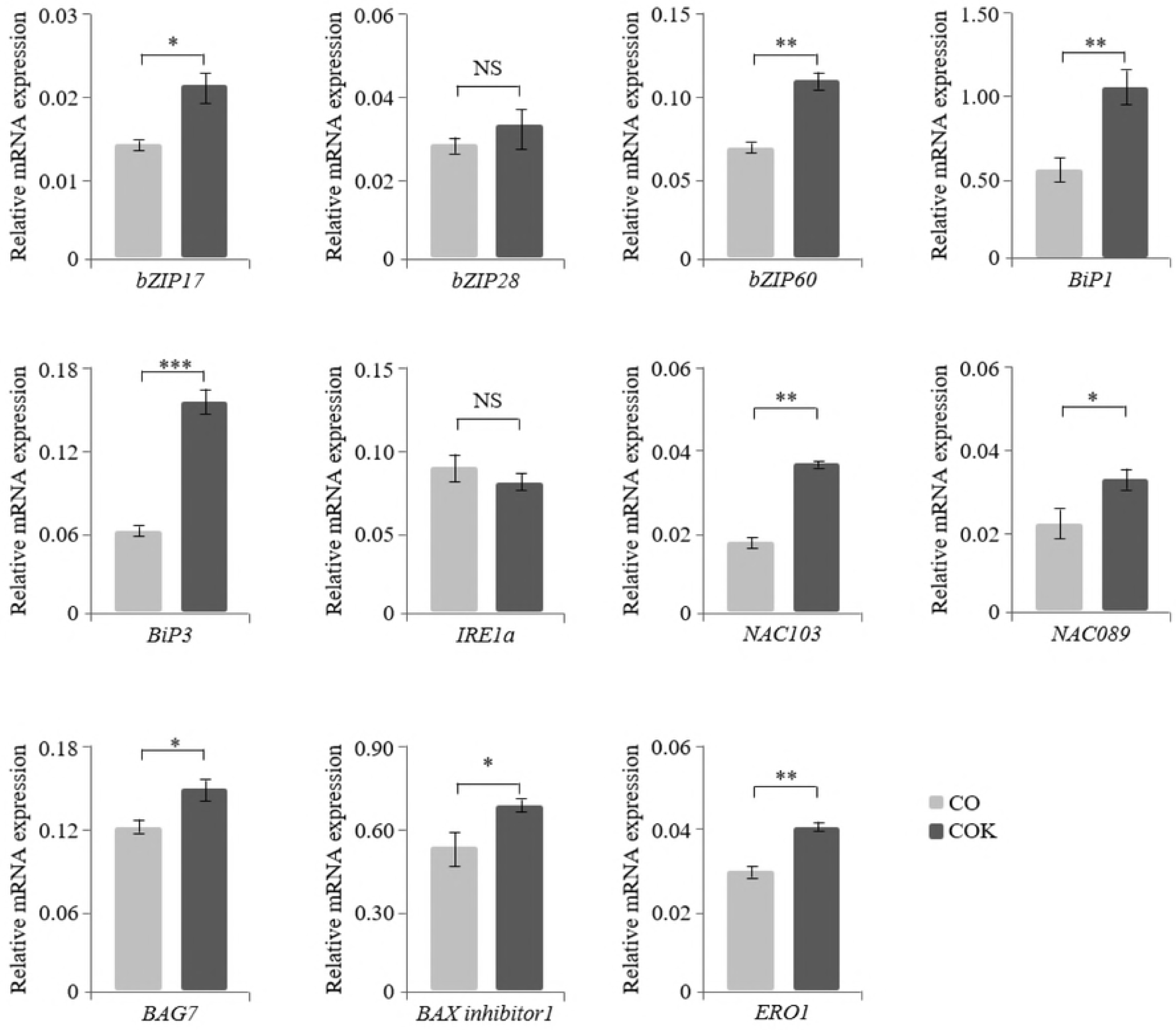
Regulation of mRNA expression of ER stress-related genes (*bZIP17, bZIP28, bZIP60, BiP1, BiP3, IRE1a, NAC103, NAC089, BAG7, BAX inhibitor 1*, and *ERO1*) in transgenic *Arabidopsis* plants expressing mAb CO and mAb COK measured with qRT-PCR analysis. RNA was isolated from the roots of plants from each experimental group (mAb CO and mAb COK) and grown on kanamycin-containing media for 12 days. The ubiquitin 10 (*UBI 10*) gene was used as a control gene for normalization. Error bars represent the mean ± standard deviation (SD) of three technical replicates with two biological replicates. Statistically significant differences are indicated by asterisks (NS: not significant; *: p < 0.05; **: p < 0.01; and ***: p < 0.001). The X-axis represents ER stress-related genes, and the relative mRNA expression levels of each transformant are expressed on the Y-axis.

### Comparisons of root lengths of CO and COK plants grown under *in vitro* and *in vivo* conditions

A total of 100 vernalized and sterilized seeds for each experimental group (CO and COK) were sown on agar plates containing kanamycin. Two-weeks after seeding on an agar plate containing kanamycin, the average root lengths of CO and COK plants were 23.2 and 12.0 mm, respectively, and the difference between the two was significant (Fig 5A,B). Following an *in vitro* antibiotic susceptibility test, true leaf-generated shoots (CO (36 shoots), and COK (32 shoots)) were transplanted to a soil pot for further study. Ten-weeks after transplanting to a pot, the average root lengths of CO and COK plants were 40.3 and 40.0 mm, respectively (Fig 5C,D). Under *in vitro* conditions, the root length of plants in the CO group was twice that of plants in the COK group. However, under *in vivo* conditions, there was no significant difference between the root length of plants in the CO and COK groups.

**Fig 5.**
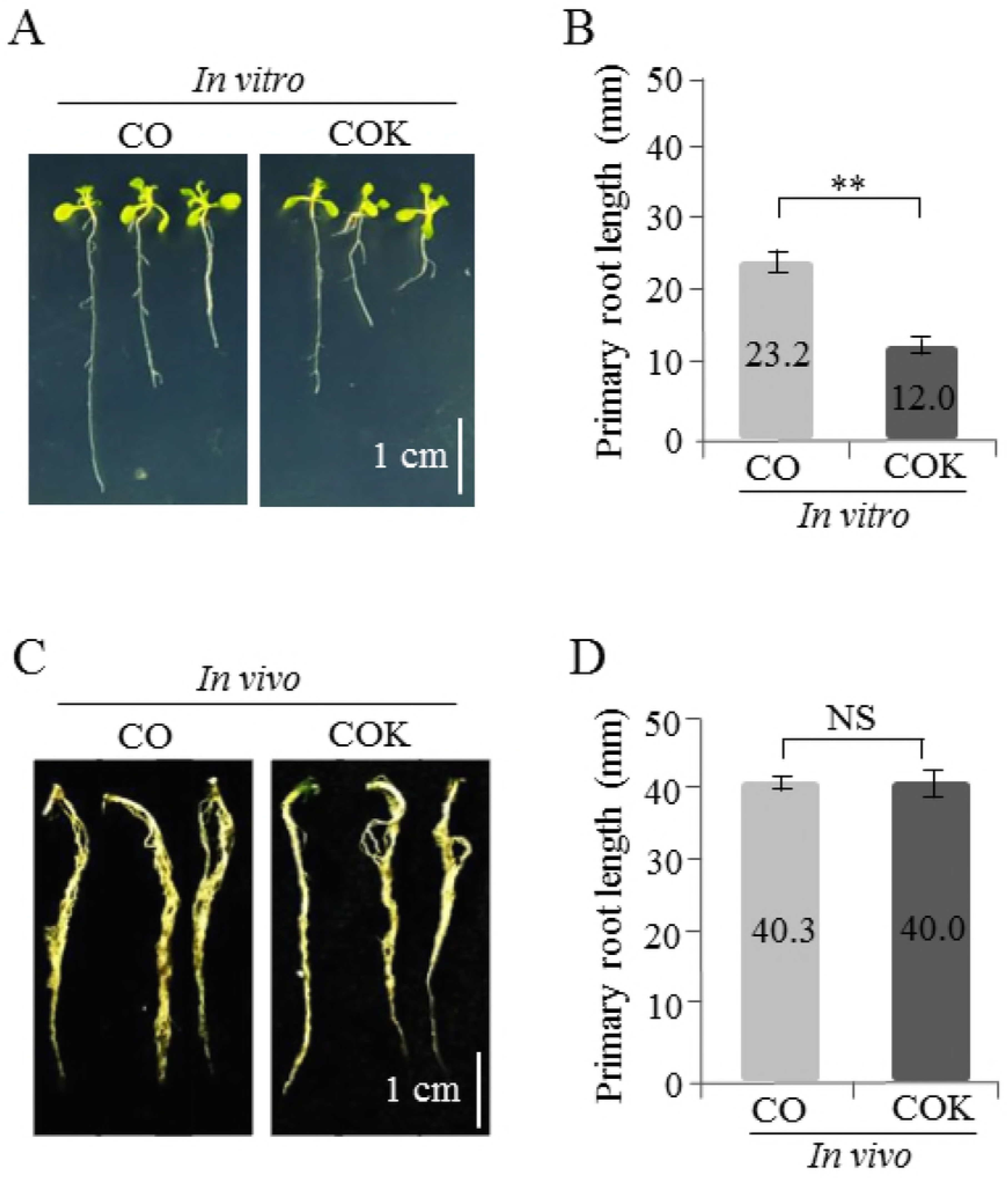
Comparison of primary root growth of transgenic *Arabidopsis* plants expressing mAb CO (CO) and mAb COK (COK) under *in vitro* and *in vivo* conditions. (A) *In vitro* germinated seedlings examined two weeks after sowing on MS media. The scale bar represents 1 cm in each photograph. (B) Comparison of average primary root lengths of transgenic *Arabidopsis* plants expressing CO and COK grown under *in vitro* conditions. (C) Roots of *in vivo* CO and COK plant seedlings 10 weeks after transplanting from *in vitro* growth to a soil pot. (D) Comparison of average root lengths of transgenic *Arabidopsis* plants expressing CO and COK grown in a chamber for 10 weeks. Error bars represent the mean ± SD. Statistically significant differences are indicated by asterisks (**: p < 0.01; NS: not significant).

### Growth trends of rosette leaves of transgenic *Arabidopsis* plants expressing mAb CO and mAb COK under *in vivo* conditions

Among the total seeds sown on kanamycin-containing MS medium, true leaf-generated shoots (NT (16 shoots), CO (32 shoots), and COK (30 shoots)) were transplanted to a soil pot for further study. One-week after transplantation to a pot, the average rosette leaf lengths (measured from the petiole to the blade) of NT, CO, and COK specimens were 0.81, 0.85, and 0.78 cm, respectively (S2B Fig). Two-weeks after transplantation to a pot, the average rosette leaf lengths of NT, CO, and COK specimens were 1.90, 1.94, and 1.88 cm, respectively (S2B Fig). Three-weeks after transplantation to a pot, the average rosette leaf lengths of NT, CO, and COK were 2.80, 2.70, and 2.65 cm, respectively (S2B Fig). Four-weeks after transplantation to a pot, the average rosette leaf lengths of NT, CO, and COK were 3.44, 3.32, and 3.22 cm, respectively (S2B Fig). Under *in vivo* conditions, the rosette leaf lengths of NT, CO, and COK were not significantly different.

### Effects of KDEL on mAbs CO and COK protein expression and localization in rosette leaf tissues

To investigate the effects of ER retention motif tagging on transcription and translation levels of mAbs CO and COK, RT-PCR and western blot analyses were conducted, respectively (Fig 6AB). Rosette leaves of the T_4_ generation of CO (30 plants, five lines), COK (25 plants, four lines), and NT (eight plants) plants were used to extract mRNA and proteins, respectively. The HC and LC transcription levels did not differ significantly between CO and COK lines. However, HC expression levels in COK plants were 4.4-fold higher than those of CO plants, and LC expression levels in COK plants were 4.9-fold higher than those of CO plants (Fig 6C). In addition, COK plants exhibited more stable expression patterns than CO plants. These results indicated that ER retention motif tagging to HC affects protein expression levels rather than transcription levels. Immunohistochemical analyses using a Dako REAL^™^ EnVision^™^ Detection System and Mayer’s hematoxylin were conducted to confirm the localization of mAb CO and mAb COK in rosette leaves of *Arabidopsis* plants (NT, CO, and COK) (Fig 6D). Brown color staining was observed in leaf tissues of both CO and COK *Arabidopsis* plants, whereas brown color stained cells were not detected in the leaf tissues of NT plants (Fig 6D). The brown color density of COK plants was stronger than that of CO plants (Fig 5D; ×;200, ×;400, and ×;1000). In COK plants, brown color staining was localized in the intracellular space, including xylem and phloem areas, whereas staining was localized in the extracellular space of CO leaf tissues (Fig 6D; ×;1000).

**Fig 6.**
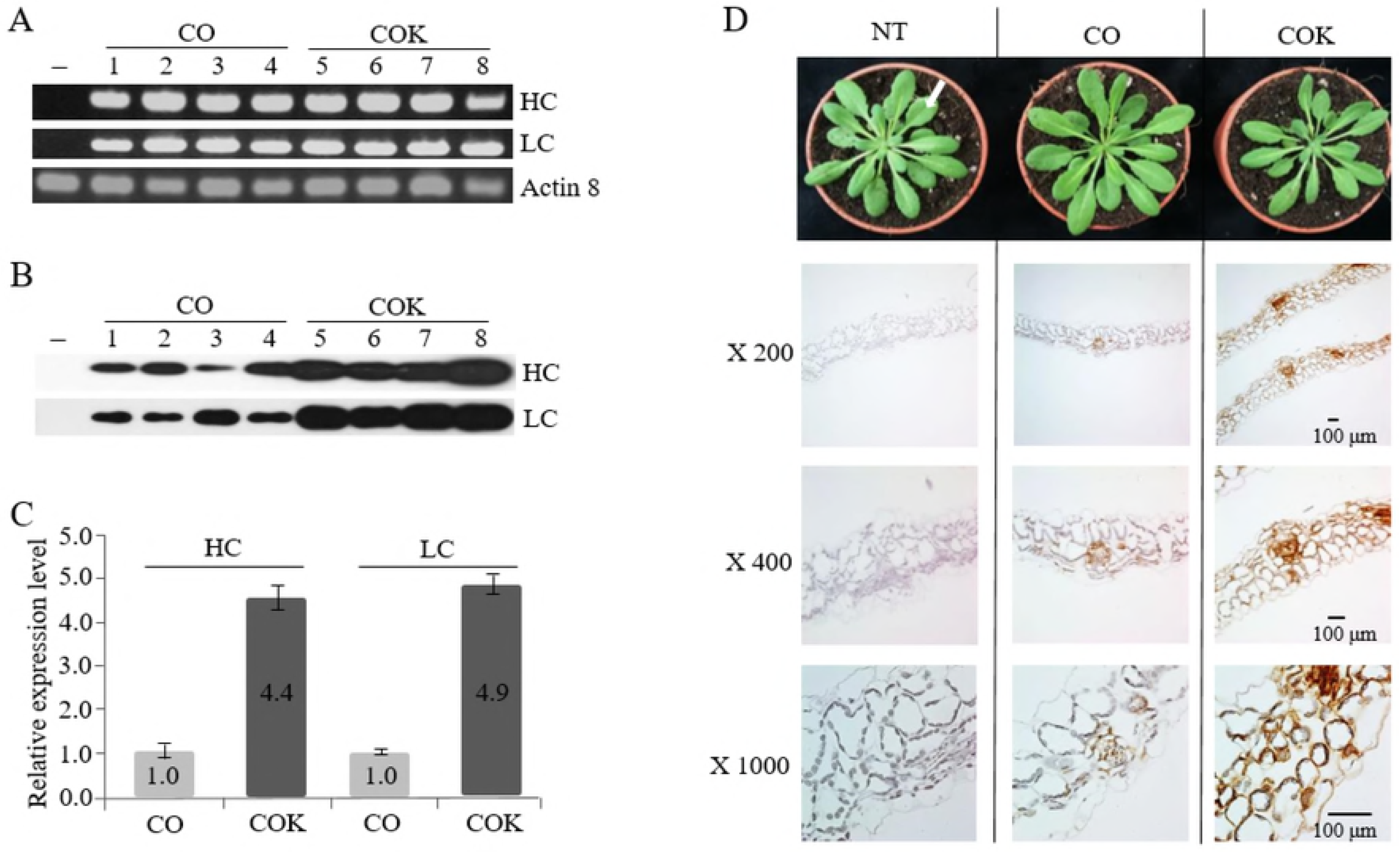
Effects of the ER retention motif on the expression and distribution of transgenic *Arabidopsis* plants expressing mAb CO and mAb COK. (A) RT-PCR and (B) western blot analyses were performed to confirm HC and LC expression based on RNA and protein expression levels. Lane 1: non-transgenic (NT); lanes 2–5: transgenic plants expressing mAb CO; and lanes 6–9: transgenic plants expressing mAb COK. (C) Relative expression levels of HC and LC determined by immunoblot analyses. Values represent the mean for each experimental group, and SD is indicated by error bars. (D) Intracellular distribution of mAb CO and mAb COK in the rosette leaf tissues of transgenic *Arabidopsis* plants. Freshly harvested rosette leaf tissues (indicated by a white arrow) were fixed in 10% formalin overnight and processed for paraffin embedding. A Dako REAL^™^ EnVision^™^ Detection System was used, according to the manufacturer’s instructions, and samples were then counterstained with Mayer’s hematoxylin. Lane 1: non-transgenic (NT) plant; lane 2: mAb CO; and lane 3: mAb COK. Slides were observed under a microscope (magnification: ×;200, ×;400, and ×;1000; BX53F; Olympus, Tokyo, Japan)]. Scale bar represents 100 μm.

### Mass production and purification of anti-CRC mAbs from *Arabidopsis* plants using protein A affinity chromatography

Amplified seeds of transgenic *Arabidopsis* CO and COK plants were sown in soil and cultivated in a greenhouse facility (Fig 7A). A total of 40 g of transgenic *Arabidopsis* leaves were harvested in each tray, and 200 g of plant biomass (each CO and COK plant) were used to purify anti-CRC mAbs (CO and COK). Ammonium sulfate-mediated total soluble protein precipitation and protein A affinity chromatography were used to purify anti-CRC mAbs from plant leaf biomass, and the two processes efficiently isolated the majority of total soluble proteins and separated specific mAbs CO and COK from plant biomass extracts, respectively. SDS-PAGE analyses were used to identify HC (50 kDa) and LC (25 kDa) of anti-CRC mAb^P^ CO17-1A from elution fractions (Fig 7B). Both mAb CO and mAb COK were mainly eluted in fraction samples (F1 and F2; Fig 7B). Each eluted sample (F1–F3) was quantified, and the total amounts of mAbs obtained were 410 (CO) and 1,950 μg (COK). The amount of mAb obtained from COK plants was five-times higher than that obtained from CO plants.

**Fig 7.**
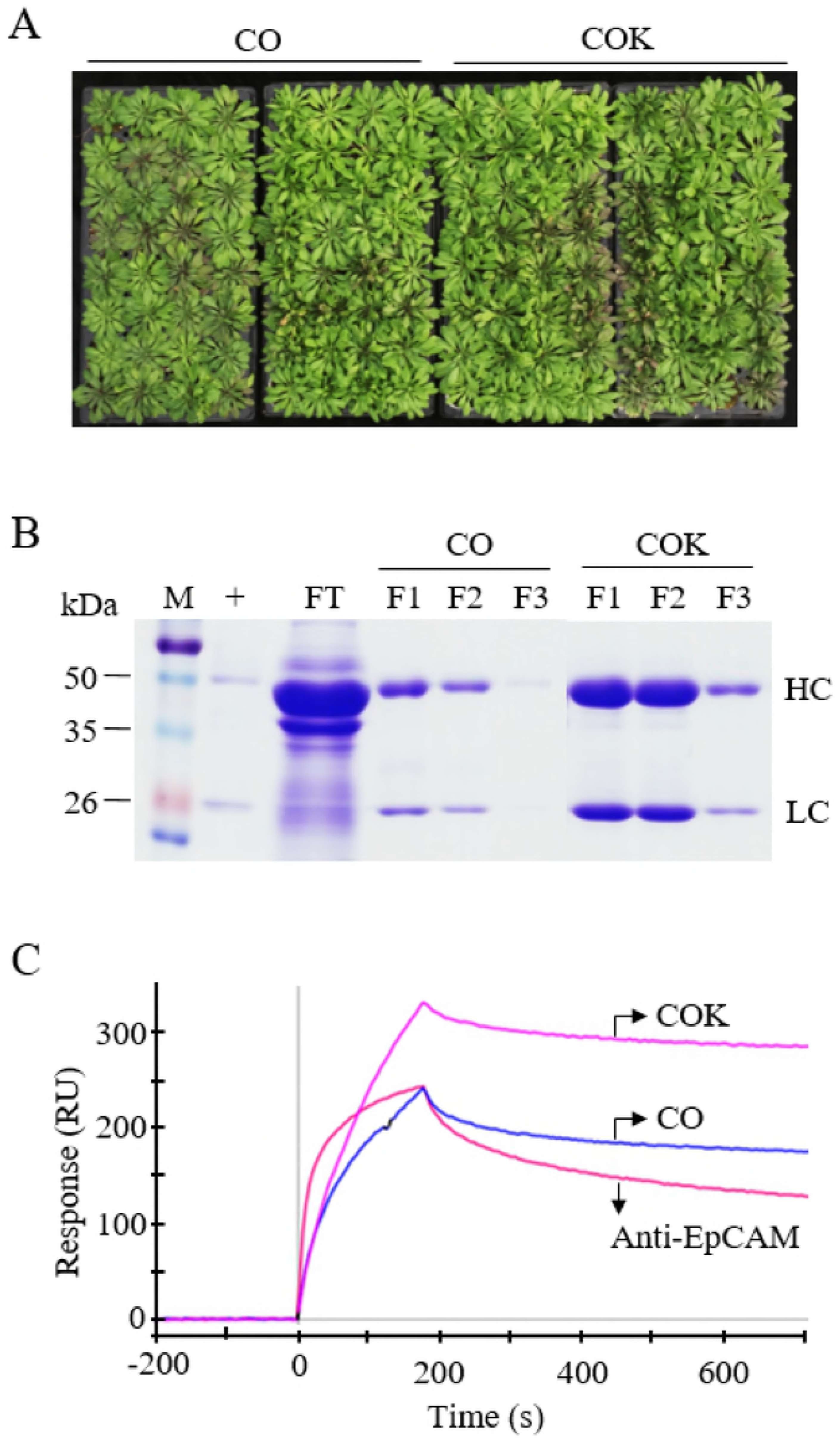
Purification and functions of mAb CO and mAb COK expressed in *Arabidopsis* plants. (A) Representative images depicting the mass production of mAb CO and mAb COK in transgenic *Arabidopsis* plants. Seeds were sown in a greenhouse and conditioned at 24°C, 30% humidity, 16 h light, and 8 h dark. Photographs were taken seven weeks after sowing homozygous seeds. (B) SDS-PAGE analysis of eluted F1–F3 fractions of purified anti-cancer mAbs obtained from *Arabidopsis* plants. Lane 1: protein marker; lane 2: anti-EpCAM mAb; lane 3: flow-through; lanes 4–6: mAb CO; and lanes 7–9: mAb COK. Samples (20 μL) were loaded into each well. (C) Surface plasmon resonance analysis was used to confirm the binding activity of mAb CO and mAb COK to an epithelial cell adhesion molecule (EpCAM-Fc). To confirm antigen-antibody binding activity of both mAb CO and mAb COK purified from transgenic *Arabidopsis* plants, the surface of a GLC sensorchip was immobilized with EpCAM fused to human IgG Fc fragment (EpCAM-Fc) molecules. Anti-EpCAM mAb (300 nM), CO (300 nM), and COK (300 nM) were applied to the sensor chip with a flow rate of 80 μL·min^−1^ at 25°C, and the response curves shown in the figure were consequently obtained.

### Binding activity of anti-CRC mAbs to antigen EpCAM proteins

Purified samples (mAb^P^ CO and mAb^P^ COK) were dialyzed with 1×; PBS and normalized to adjust the concentration (25 μg·mL^−1^) for SPR analyses (Fig 7C), and 250-μL samples (anti-EpCAM mAb, mAb^P^ CO, and mAb^P^ COK) were used for each SPR analysis. Both mAb types (mAb CO and mAb COK) had higher and similar binding affinities to EpCAM compared to mammalian-derived anti-EpCAM mAb. Results of SPR analyses showed that COK had the highest equilibrium concentration to the immobilized chip coated with EpCAM-Fc after 200 seconds (Fig 7C). Anti-EpCAM mAb and mAb CO had lower equilibrium concentrations compared to mAb COK, and dissociation rates were almost equal between CO and COK. However, anti-EpCAM mAb dissociated faster than both CO and COK (Fig 7C).

## Discussion

This study demonstrated the effects of ER retention motif KDEL on plant growth phenotypes and anti-CRC mAb expression in transgenic *Arabidopsis* plants. The effects of KDEL tagging were previously discussed in transgenic tobacco plants, and it resulted in the retention of KDEL-tagged recombinant proteins in the ER, leading to a high mannose glycan protein structure [25, 26]. In the current study, *A. thaliana*, used for molecular biofarming, was used to express mAb CO and mAb COK.

T_1_ seeds were obtained using a traditional floral dip *Arabidopsis* transformation protocol, and positive transformants were screened by PCR and western blot analyses, consequently leading to the identification of stable homozygous transgenic lines. Sterilized seeds of transgenic *Arabidopsis* plants, expressing mAb CO and mAb COK, and NT plant seeds were vertically sown on kanamycin or non-kanamycin media to confirm the effects of ER retention motif tagging on germination rates and the primary root growth of each group. mAb CO or mAb COK transgene insertions in *Arabidopsis* plants had little influence on germination rates, and this result was similar to that of the NT group. However, the main root lengths of plants expressing mAb COK were approximately two times shorter than those of plants expressing mAb CO. We hypothesized that the accumulation of anti-CRC mAbs in the ER might induce ER stress-related genes in *Arabidopsis* plants, consequently affecting the primary root phenotype of transgenic *Arabidopsis* plants [27, 28]. Previous studies demonstrated that the accumulation of recombinant proteins in the ER might induce ER stress, which activates UPR to reduce ER stress and maintain homeostasis [8, 9, 28]. Among the three major UPR branches [12], *ATF6* and *IRE1* pathway-related genes were analyzed in the present study. The main transcription factors related to the UPR pathway (*bZIP17, bZIP60, BiP1, BiP3, NAC103, NAC089, BAG7, BAX inhibitor 1*, and *ERO1*) [27] were upregulated in the roots of plants expressing mAb COK compared with those of plants expressing mAb CO.

*In vitro* seedlings that survived antibiotic selection were transferred to soil pots in a growth chamber to confirm plant growth, transcription levels, and translation levels of mAbs in each experimental group. Unlike *in vitro* conditions, plant phenotypes such as rosette leaf and primary root length of mAb COK plants were similar to those of the mAb CO and NT groups under *in vivo* conditions. In addition, genes related to ER stress and UPR did not differ significantly between mAb CO and mAb COK expressing plants grown under *in vivo* conditions (data not shown). It is speculated that conditions such as light, fertilizer, and soil (non-kanamycin added) led to reduce stress compared to *in vitro* conditions.

Although C-terminal KDEL tagging induced ER stress and resulted in slightly altered plant growth phenotypes, the ER retention motif boosted the amount of translated mAb COK per unit to nearly four times higher in fresh leaves, and it exhibited a more stable expression pattern compared to mAb CO [29, 30]. The transcript levels of mAb HC and LC genes were nearly equal between mAb CO and mAb COK plants, and these results suggested that ER retention motifs efficiently accumulated recombinant anti-CRC mAbs in plant tissues.

The protein expression-boosting effects of KDEL tagging were confirmed by immunohistochemical analyses using fresh rosette leaf tissues obtained from transgenic *Arabidopsis* plants expressing mAb CO and mAb COK. Our study showed that ER retention motif KDEL led to mAb COK being well-retained in the intracellular space without any damage to plant cells, consequently inducing high protein accumulation levels per unit leaf biomass.

To obtain anti-CRC mAbs, approximately 2,000 seeds from each group (mAb CO and mAb COK) were sown in a greenhouse facility, and mature plants were harvested for purification. No difference was observed in the phenotypes of transgenic plants, and this was consistent with growth chamber culture results. Both mAb CO and mAb COK expressed from transgenic *Arabidopsis* plants were efficiently purified with ammonium sulfate-mediated precipitation and protein A affinity column processes, without any deviation from the original protocol [31]. SDS-PAGE results showed that overall amounts of mAb^P^ CO per unit mass were greatly increased up to five times in mAb COK plants compared to mAb CO plants.

The binding affinities of both mAb^P^ CO and mAb^P^ COK to target antigen EpCAM molecules were higher or similar to mammalian-derived anti-EpCAM mAb counterparts, respectively. Moreover, these results were previously discussed in transgenic tobacco system [18]. Taken together, our findings suggest that *Arabidopsis* plants are a promising platform for the generation of anti-CRC mAbs with biological activities that are consistent with the anti-EpCAM mAb, a counterpart antibody. The introduction of the ER retention motif enabled the bulk production of mAbs in plants with enhanced ER accumulation, without the generation of significant *in vivo* stress regardless of *in vitro* ER stress. Therefore, an ER retention strategy is recommended to induce high accumulation of recombinant antibodies in a plant expression system.

## Acknowledgments

This research was supported by a grant (Code# PJ0134372018) from the Korean Rural Development Administration and the National Research Foundation of Korea Grant that was funded by the Korean Government (MEST) (NRF-2017R1A2A2A0569788).

## Author contributions

KK, IC, and YJK contributed to the planning and design of the experiments. YKL supplied materials and analysis tools and performed qRT-PCR analyses. IC manufactured paraffin blocks with plant leaf tissues and performed pathological experiments. KK, IC, and S-CM analyzed and interpreted all data, and they wrote the manuscript. All authors checked the concept of this study and carefully confirmed the original data of this manuscript before submission.

---

**Fig S1.**
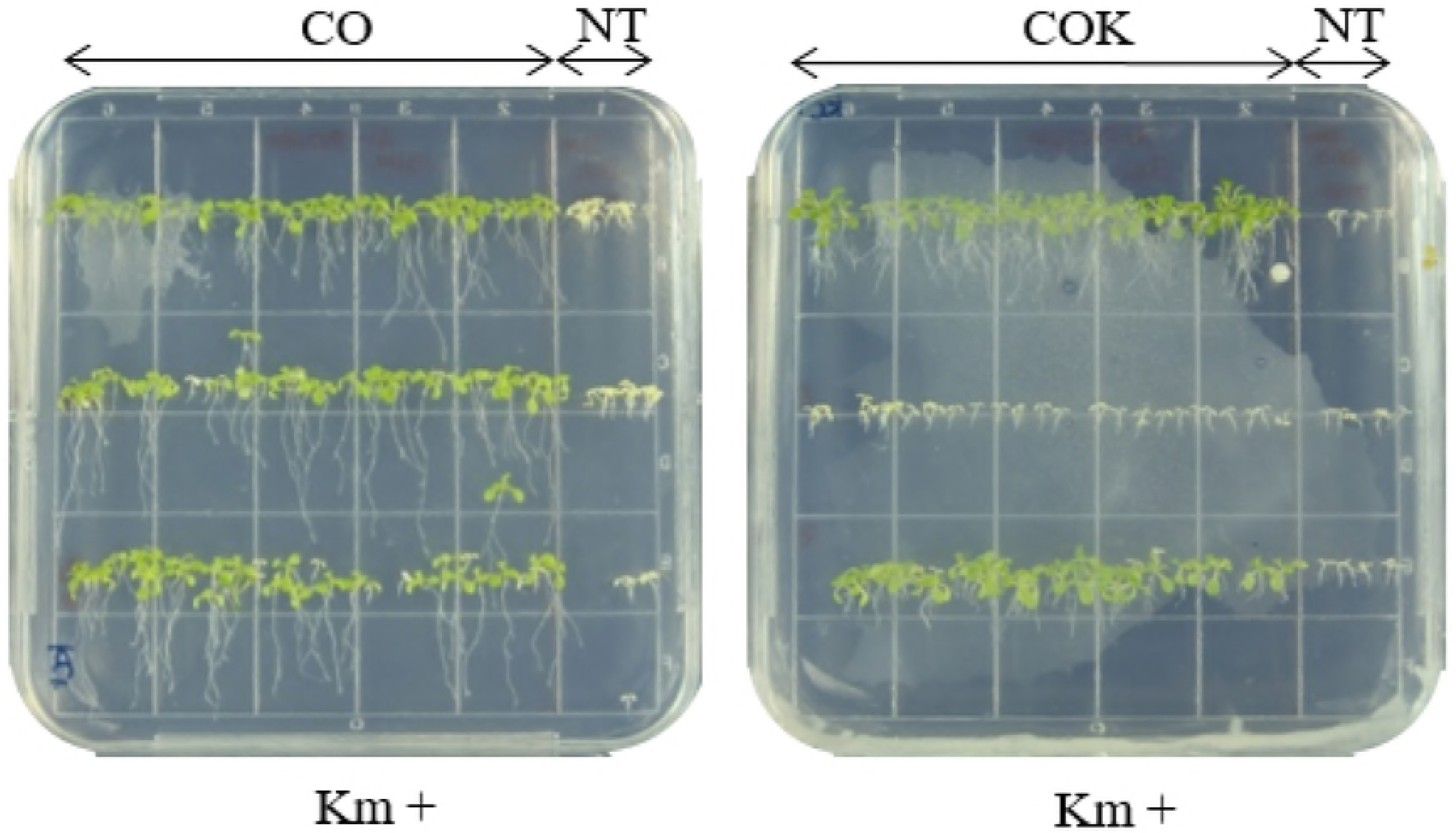
Selection of homozygous lines of transgenic *Arabidopsis* plants expressing mAb CO and mAb COK. Kanamycin selection was repeated over generations to obtain homozygous seeds for further study. Photographs were taken 14 days after seed germination using a camera (Digital Gross System) (Humintec, Suwon, Korea). CO: seedling expressing mAb^P^ CO17-1A; COK: seedling expressing mAb^P^ CO17-1AK; Km+: kanamycin containing media.

**Fig S2.**
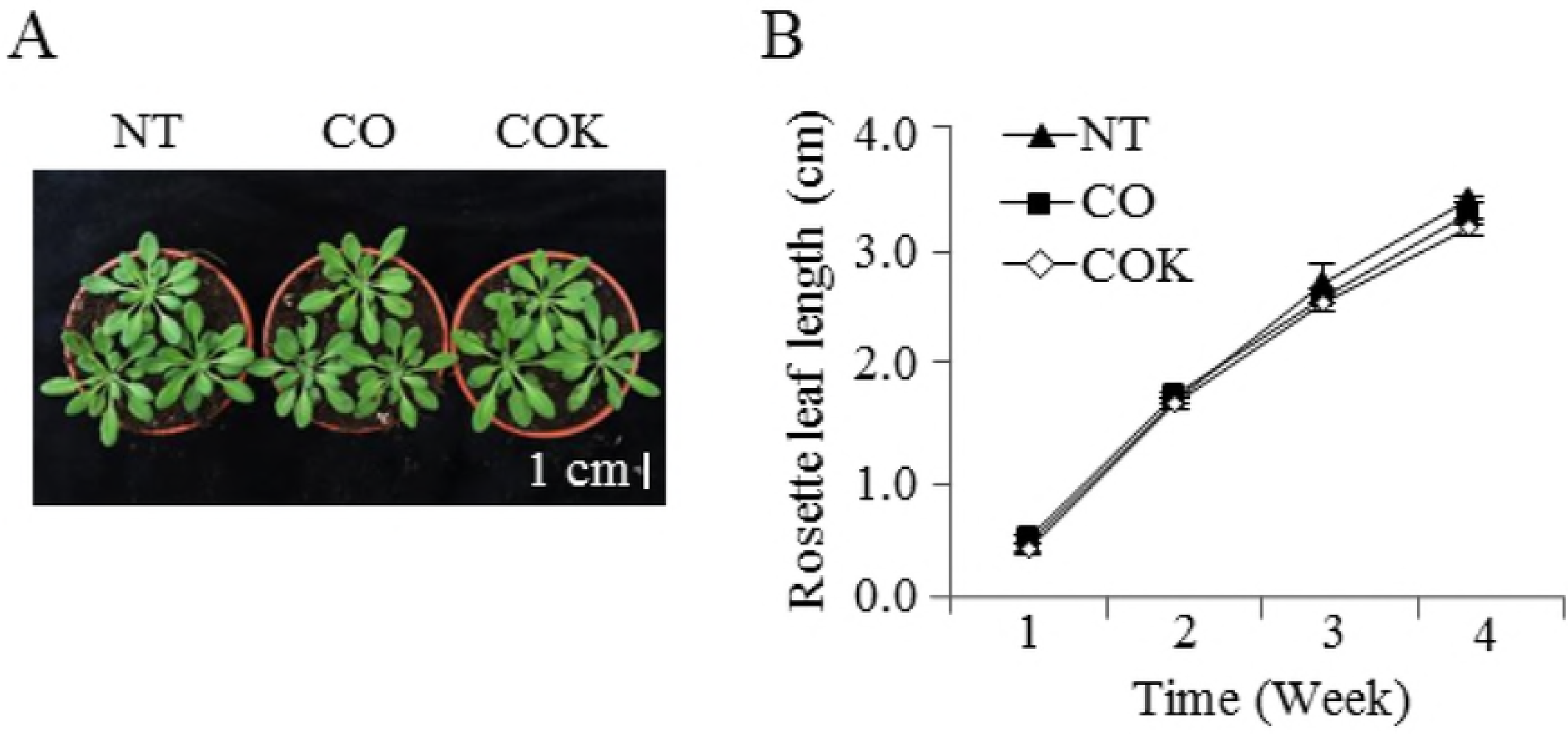
Comparison of rosette leaf lengths between transgenic *Arabidopsis* plants expressing CO and COK. (A) Seeds (NT, mAb CO, and mAb COK) were sown into pots containing soil in a growth chamber. Plants were photographed one month after transfer to pots. NT: non-transgenic plants; CO: transgenic plants expressing mAb CO; COK: transgenic plants expressing mAb COK. (B) Rosette leaf lengths were measured from the petiole to the blade using a ruler at 1-week intervals after transplantation from *in vitro* conditions. Scale bar represents 1 cm in each photograph.

